# A new platform for high-throughput therapy testing on iPSC-derived, immature airway from Cystic Fibrosis Patients

**DOI:** 10.1101/2021.07.05.451013

**Authors:** Jia Xin Jiang, Leigh Wellhauser, Onofrio Laselva, Irina Utkina, Zoltan Bozoky, Tarini Gunawardena, Zoe Ngan, Sunny Xia, Paul D.W. Eckford, Felix Ratjen, Theo J. Moraes, John Parkinson, Amy P. Wong, Christine E. Bear

## Abstract

Induced pluripotent, stem cell (iPSC)-derived models of airway tissue have successfully modeled the primary defect in regulated chloride conductance caused by the major Cystic Fibrosis causing mutation, F508del. However, it remains unclear if iPSC-derived airway cultures can be used in high-throughput therapy development for F508del and rarer mutations. There is an urgent need for airway tissue models that reflect the variability of patient-specific responses and are scalable for therapy development. In the current work, we describe a robust, high-throughput fluorescence assay of mutant CFTR function in iPSCs differentiated to immature airway epithelium. This assay measures reproducible functional responses to modulators targeting either the major CF mutant F508del or the nonsense mutant: W1282X-CFTR. We show that the ranking of patient-specific responses to interventions in this stem-cell based model recapitulates the ranking observed in primary nasal epithelial cultures obtained from the same individuals. In summary, these proof-of-concept studies show that this novel platform has the potential to support therapy development and precision medicine for Cystic Fibrosis patients.

**One Sentence Summary:** We describe a fluorescence-based platform that enables high-throughput Cystic Fibrosis therapy testing using iPSCs differentiated to immature lung.

## INTRODUCTION

Mutations in the *CFTR* gene can result in the disease called Cystic Fibrosis (CF). The major mutation, called F508del leads to CFTR protein misassembly, misprocessing, mistrafficking and altered function as a phosphorylation regulated chloride channel (*1–3*). Combinations of small molecule modulators called correctors, that rescue misassembly, together with potentiators that increase the probability of channel opening have been shown to be effective in improving lung function in individuals harboring the major mutation, F508del (*4–7*). However, not all individuals with this mutation show the same level of clinical improvement, supporting the ongoing need for novel therapy development. In addition, there are multiple, rarer CF-causing mutations for which there are no approved therapies. For example, W1282X causes the loss of CFTR mRNA expression due to the triggering of nonsense mediated decay (NMD), a mechanism thought to vary amongst individuals (*8*). The process of NMD hampers therapies promoting read-through of premature stop codons and in the case of W1282X, protein modulators known to rescue the altered functional expression of the shorter protein (*9–11*). Therefore, there is an urgent need for novel therapy development for the treatment of Cystic Fibrosis. Tissue models that reflect the variability amongst patient-specific responses and are scalable to high-throughput formats would facilitate such therapy development.

Traditionally, in-vitro studies of CFTR modulators, have employed bronchial epithelial tissue cultures from explanted CF lungs obtained at the time of transplant. Such “gold-standard” studies were, and continue to be, very instructive with respect to understanding the mechanism of modulator activity and evaluating their efficacy in relevant tissues. The functional consequences of modulator treatment are examined in direct measurements of ion conductance in the Ussing chamber apparatus (*12–14*). Although informative, these methods are low-throughput and not readily conducive to comparative studies of multiple interventions simultaneously. In addition, as the cell donor has undergone a lung transplant, the potential for directing individualized therapy is limited. Further, for rare mutations, for which currently no effective therapies are available, reduced availability of explanted tissue harboring the appropriate genotype impedes drug development.

In lieu of bronchial epithelial cells, the development of systems for the culturing of the nasal epithelium and rectal organoids, greatly enhanced access to patient specific tissues harboring different disease-causing mutations (*15–22*). For example, Ussing chamber studies of patient specific nasal epithelial cultures have been used to show the relative efficacy of the new modulator combination TRIKAFTA™ in rescuing F508del-CFTR and certain rare mutations (*23, 24*). In addition, the potential efficacy of novel compounds from academic groups or different pharmaceutical companies have also been revealed in Ussing chamber studies of patient-derived primary nasal epithelial cultures. Interestingly, our group and others observed that patients with the same genotype can exhibit different in-vitro drug responses (*11, 25–30*). The molecular basis for interpatient variations in in-vitro response size remains unknown but is thought to be a harbinger of the extent of variability in clinical response size (*15*).

Thus, there is a need to develop a higher-throughput testing platform of patient-specific airway tissue that enables therapy development, particularly for those CF-causing mutations like W1282X for which no therapies exist. Unfortunately, nasal epithelial cell cultures have a limited ability to expand in culture and lose CFTR functional expression with progressive population doublings (*16*). These properties are limiting to combinatorial studies of investigational compounds. Robust protocols have been developed for the differentiation of CF patient derived iPSCs (*31–37*) thereby providing the potential for a renewable source of patient derived tissue that is scalable to high-throughput testing. In proof of concept studies, the functional rescue of F508del-CFTR chloride channel activity by CFTR modulators was shown for lung tissue differentiated at the air-liquid interface (ALI) using either the Ussing chamber or a membrane potential sensitive dye (FLiPR) (*35, 38*). However, the protocol for differentiating mature airway at ALI is lengthy and not readily scalable. In the current project, we asked if a truncated protocol, generating immature airway tissue from patient-derived iPSCs could be used for drug profiling in a higher throughput format. We show here that this abbreviated differentiation protocol generated relevant CFTR expressing cell types, enabled comparative studies of multiple interventions targeting either F508del or W1282X-CFTR and recapitulated patient specific responses observed in gold-standard Ussing chamber studies of primary nasal epithelial cultures in proof-of-concept studies. We anticipate that these findings will enable the use of patient-specific iPSCs for CF precision medicine approaches.

## RESULTS

### Differentiation of IPS cells to immature lung in high-throughput format

We employed the protocol developed by Wong et al (*35*), **Fig. 1A**) to differentiate non-CF and CF iPS cells to immature lung tissue in a 96 well plate format. These cultures remained submerged under differentiation media hence, we refer to them as “submerged cultures”. As expected, these cultures express NKX2.1, a marker of lung progenitors and pan-cytokeratin – an epithelial cell marker, (**Fig. 1B**).

**Fig. 1:**
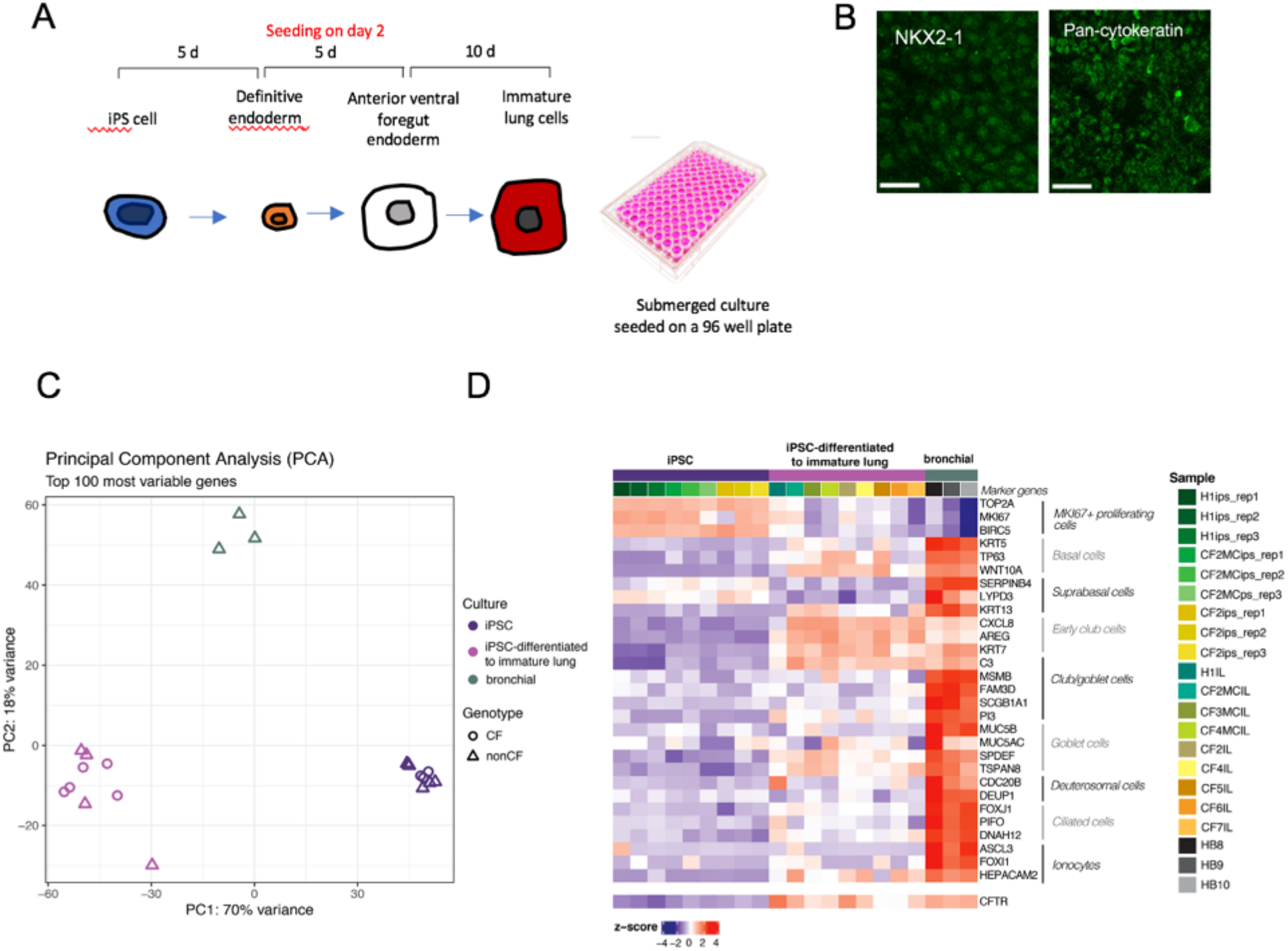
Differentiation of patient derived iPS cells to immature airway. (**A**) Schematic of differentiation protocol and timeline: human iPSCs were directed to definitive endoderm and passaged onto 96 well plates during the anterior foregut endoderm differentiation. The submerged cultures were differentiated for 10 more days into immature lung cells. (**B**) Immunofluorescence images of submerged, immature cultures. Scale bar, 20 μm. Most cells stained positive for TTF1 (NKX2-1) and pan-cytokeratin. (**C**) Principal component analysis (PCA) comparing iPS cell lines, immature lung cultures differentiated from iPS cell lines and primary bronchial cultures. Both CF and non-CF (including mutation corrected) iPS cell lines studied. (**D**) Heatmap of gene expression clustered according to cell types using marker genes (*39*). The columns correspond to different donors and whether lines are CF or non-CF (these include mutation corrected (MC). Supplementary Table 1, provides a legend for colours used with CFTR genotype, for each of the samples. Columns are also clustered as iPS cell lines, immature lung (IM) differentiated from iPS cell lines and primary bronchial cultures. Relative CFTR expression across cultures is shown in the bottom row of the heat map.

We conducted transcriptomic analyses in order to further characterize the properties of submerged cultures. Bulk RNAseq was performed for: 1) 3 undifferentiated iPS cells, each with 3 replicate cultures; 2) 9 iPS cell lines differentiated under “submerged” conditions to create immature lung cultures (4 non-CF and 5 CF); and non-CF differentiated bronchial airway cultures (n=3) as positive controls (NIH Tissue Core- University of Iowa), (**Fig. 1C and D**). Principal component analyses showed that the immature lung cultures were intermediate (as determined by PC1) between the iPS cells and differentiated mature bronchial epithelium with respect to gene expression of the 100 genes displaying the greatest variability across samples (**Fig. 1C**). Further, at least with these samples, CFTR genotype (with or without a CF causing mutation), did not change the distribution of cell sub-populations across the different samples. The heatmap in **Fig. 1D**, confirms that the immature lung cultures express genes specific for multiple airway cell types, including basal cells, early club cells and goblet cells (*39*). Interestingly, after differentiation to immature lung cultures, CFTR is expressed, close to levels comparable to those measured in mature non-CF bronchial cultures differentiated at the air-liquid interface.

### Immature lung tissue differentiated from iPSCs express functional CFTR

Cyclic AMP activated Wt-CFTR channel function was detected for immature airway cultures differentiated in wells of a 96 well plate, using a fluorescence-based assay of membrane potential changes (FLiPR) (*40*). The trace in **Fig. 2A**, shows representative forskolin mediated membrane depolarization in the presence of an outward chloride gradient (mean change −/+SEM for 4 wells). This response was inhibited by the CFTR inhibitor, CFTRInh-172 as expected for CFTR mediated channel activity. Expression of the mature CFTR protein as a 180 kD polypeptide was confirmed for cultures grown in this format (**Fig. 2B**).

**Fig. 2:**
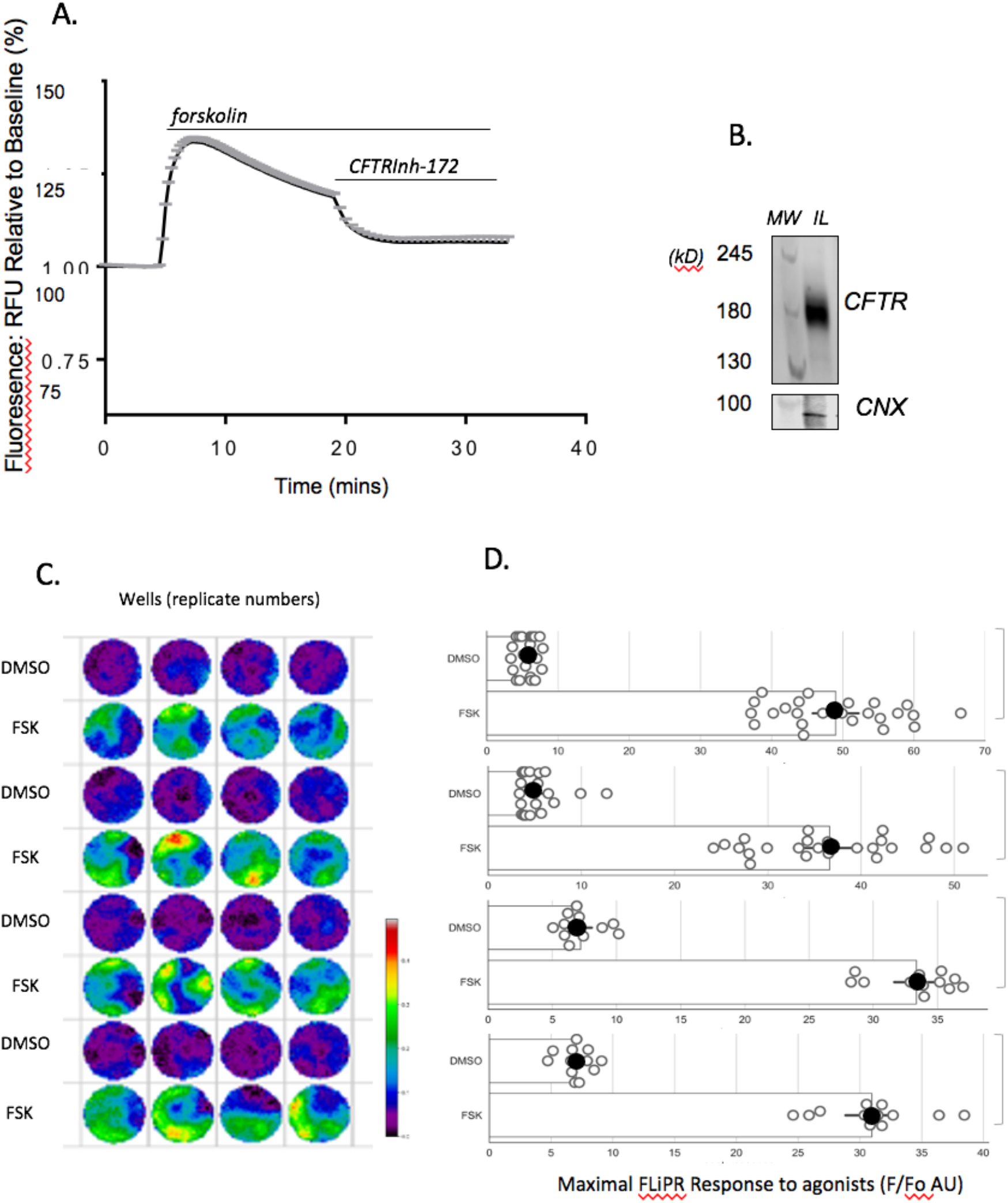
Immature lung cultures express Wt CFTR and channel activity in fluorescence-based assay. (**A**) Representative FLiPR trace (mean−/+SEM) responses of 4 wells plated with immature airway cultures stimulated by 10 μM forskolin and inhibited by 10 μM CFTR Inh-172. This cell line was derived from donor=CF2MC for which the F508del mutation was corrected to Wt. The naming of lines is consistent with Figure 1. (**B**) Western blot shows MW markers and mature CFTR expression (180 kD). Calnexin (CNX) was used as loading control. (**C**) Sample study for establishing assay statistics (Z-factor and SSMD). Multiple wells seeded with iPSC differentiated to immature airway epithelium. Alternating rows (each containing 4 replicates) show responses to agonist (FSK) or vehicle control (DMSO). The response size is colour-coded as shown in the scale bar, red showing the highest response. **(D)** Bar graph shows the peak responses of Wt CFTR function (after vehicle (DMSO) or agonist (FSK) in each well (open circle) of a 96 well plate. Bar graphs summarize data from four, 96 well plates generated from 2 differentiations from the iPSC line, CF2MC. Data from each plate grouped with a bracket. The solid symbol represents the mean for each condition in a plate.

The heat map in **Figure 2C**, shows well scans of FLiPR dye fluorescent change with depolarization caused by forskolin or the DMSO vehicle added to alternating rows in quadruplicate. Red shows the highest responses and dark blue, the lowest responses. Interestingly, for certain wells, there are focal hot spots of activity, reflecting heterogeneity of the culture. The open circles in the bar graph of **Fig.2D**, show the mean peak CFTR channel activity after DMSO or forskolin measured by FliPR for each well in a 96 well plate. The solid symbol represents the mean of all the technical replicates for either DMSO or forskolin on a plate. This FLiPR assay of CFTR mediated chloride conductance in immature airway cultures exhibits reproducibility in a 96 well plate format with a Z-factor score of 0.33 and an SSMD (Strictly standardized mean difference) score of 4.59. These metrics support the claim that this is a good to excellent assay platform for CFTR channel function (*41, 42*).

### Immature lung tissue differentiated from CF iPSCs models primary defects caused by F508del

Having shown that the FLiPR assay of immature airway differentiated from iPSCs is suitable for a high-throughput assay of Wt-CFTR, we applied it to study of the mutation, F508del-CFTR, and its modulation by small molecules. In Fig. 3, we show that the primary defects caused by the major mutation, F508del are recapitulated in immature lung cultures generated from two different patient donors who are homozygous for this mutation. The donors identification numbers are consistent those shown in Fig.1. For immature airway cultures from both CF donors: CF2 and CF4, the mean residual forskolin activated CFTR channel activities, (5.67+/−2.16 (SD), n=4, and 7.25+/−0.91 (SD), n=4) were significantly reduced relative to that exhibited by non-CF culture, (37.50+/− 8.02 (SD, n=4) as shown in Figs 3 and 2 respectively. As expected, the abundance of mature band C was also reduced for the major mutation in iPSC-derived airway epithelium in the absence of small molecule modulators (Fig. S1).

**Fig. 3:**
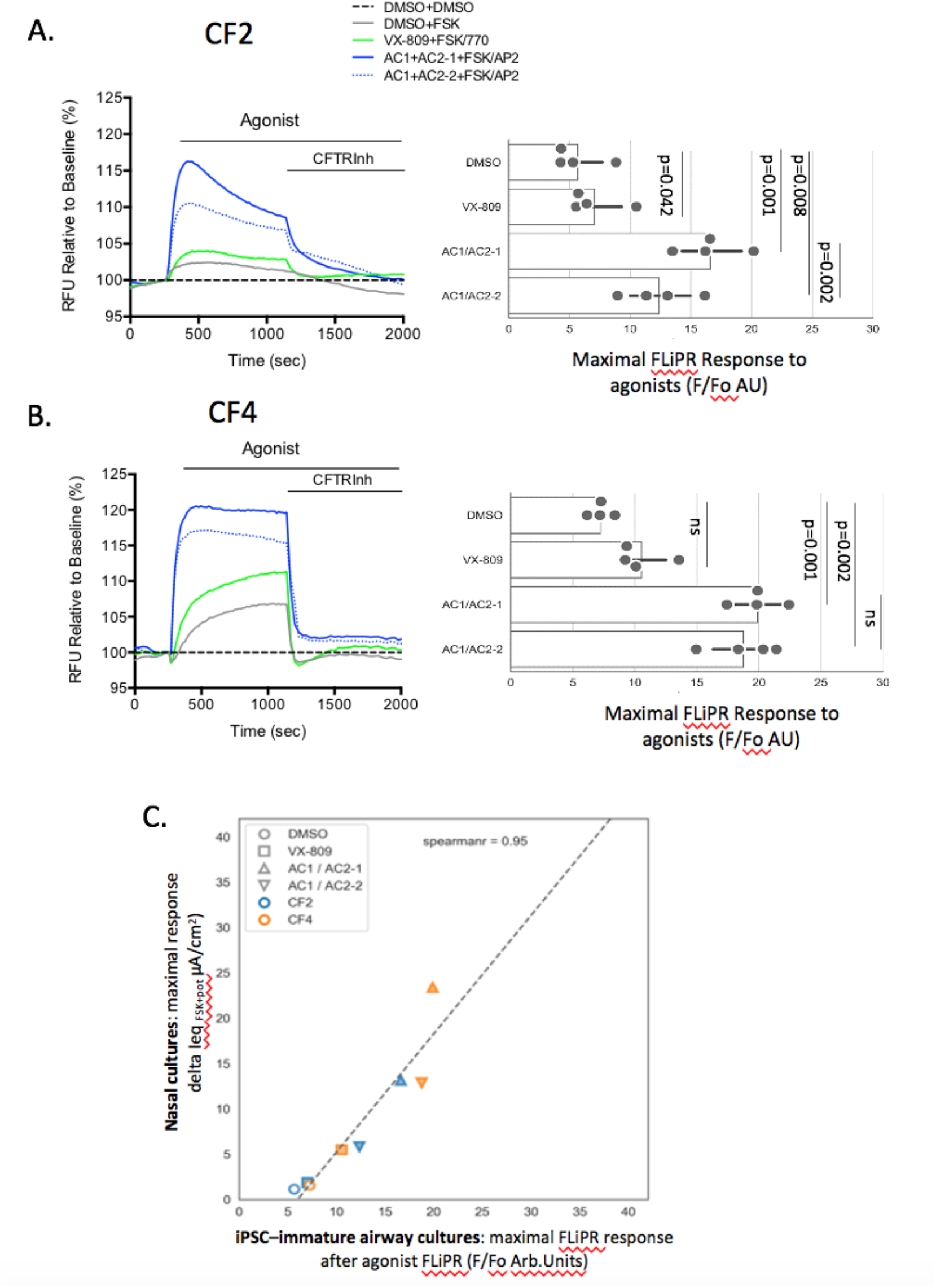
Immature lung cultures generated from iPSCs from two different donors homozygous for F508del exhibit robust, responses to modulators. (**A)** Left panel: Representative FLiPR traces of cultures derived from iPSCs with F508del mutation after chronic rescue (48 hours) with DMSO (0.1%) or small molecules as defined in the legend with concentrations indicated in **Table 1**. After 5 min baseline, the cells were stimulated with DMSO or FSK +/− 1μM VX-770 (or 1.5 μM AP2). Right panel: Bar graphs show peak responses from each modulator combination. Each solid circle represents mean peak responses of 4 wells in a 96 well plate for 4 plates of immature airway epithelium generated from a single differentiation of iPSC line: CF2. The naming of lines is consistent with Figure 1. **(B)** Left: FLIPR traces showing responses to small molecule modulators on immature airway epithelium differentiated from an iPSC line derived from donor: CF4. Right: Bars show reproducibility of FLIPR assay. Each solid dot represents the mean peak response for 4 technical replicates (wells) of a 96 well plate and there were 4 plates generated from a single differentiation of iPSC line, CF4. (**C**) Correlation between mean donor specific activations measured using FLiPR (mean values, −/+ interventions, extracted from bars above) and mean, donor specific changes measured in the Ussing chamber, delta Ieq (μA/cm^2^) after forskolin and treatment as reported in (*44*).

Treatment of immature lung cultures from donor, CF2, with small molecule modulator combinations led to reproducible responses in the FLiPR assay **(Fig. 3A)**. Relative to the vehicle (DMSO) control, treatment with the corrector, lumacaftor, i.e., VX-809, led to a partial rescue of the functional expression for CF2 immature lung cultures relative to vehicle (DMSO) in the presence of the potentiator (VX-770). As expected, there were robust functional responses to investigational modulator combinations: AC1+AC2-1 or AC1+AC2-2 and the potentiator: AP2 using the FLIPR assay on immature lung cultures (Fig. 3A). The efficacy of these modulator combinations was published previously for studies on primary nasal epithelial cultures (*20, 25, 43*).

We compared modulator responses immature lung cultures generated from donor, CF4, another individual who is homozygous for F508del. Whilst the ranking of modulator efficacies was similar for the two donors; CF2 and CF4, the responses to the modulator combination: AC1+AC2-2 was higher in cultures from donor CF4 than in donor CF2 (p=0.022). Together, these findings suggest that donor-specific differences in in-vitro response to modulators can be measured using this platform.

In **Fig. 3C**, we show that the responses to modulators observed for immature lung cultures generated from CF2 and CF4, correlate (r=0.95) with the *in-vitro* responses previously measured for well differentiated primary nasal cultures from the same donors (*44*). The bioelectric responses to modulators were previously measured for primary nasal cultures in Ussing chambers, considered the “gold standard” for measuring CFTR channel function. Interestingly, as in the FLiPR assay of immature lung, the fully differentiated nasal epithelial cultures generated from donor CF4 consistently exhibited as higher functional response to AC1-+AC2+2 than the nasal cultures generated from donor CF2 (p=0.022). Hence the patient-specific responses to modulators of the major mutation (F508del) observed in the high-throughput, FLiPR based assay recapitulated those observed in the gold-standard, but, low-through put Ussing chamber assay.

### Immature lung tissue differentiated from CF iPSCs models primary defects caused by W1282X and report differential responses to investigational rescue compounds

Having shown that the FLiPR assay of immature lung cultures differentiated from iPSC lines faithfully recapitulates therapeutic responses to modulator drugs targeting the major mutation: F508del, we then tested the fidelity with which this novel platform can used to measure patient-specific responses to investigational compounds targeting the rare, nonsense mutation: W1282X. The format for these phenotypic assays is shown in **Fig 4A**. Here, we show a heat map for the FLiPR assay of submerged lung cultures. Wells (24) seeded with submerged lung containing Wt-CFTR (CF2 mutation corrected (MC), are displayed on the left. Wells (32) seeded with immature lung differentiated from iPSCs generated from a donor who is homozygous for the rare nonsense mutation: W1282X (CF7) are shown on the right. The colour scale on the right of the wells corresponds to the peak FLiPR fluorescence after activation and potentiation of CFTR channel activity. The wells seeded with non-CF (Wt), immature lung tissue exhibit robust activation by forskolin and the peak corresponds to red. On the other hand, the colour of the wells containing immature lung cultures from a donor homozygous for the nonsense mutation varied (from dark blue to red, reflecting low to high responses) according to the treatments provided.

**Fig. 4:**
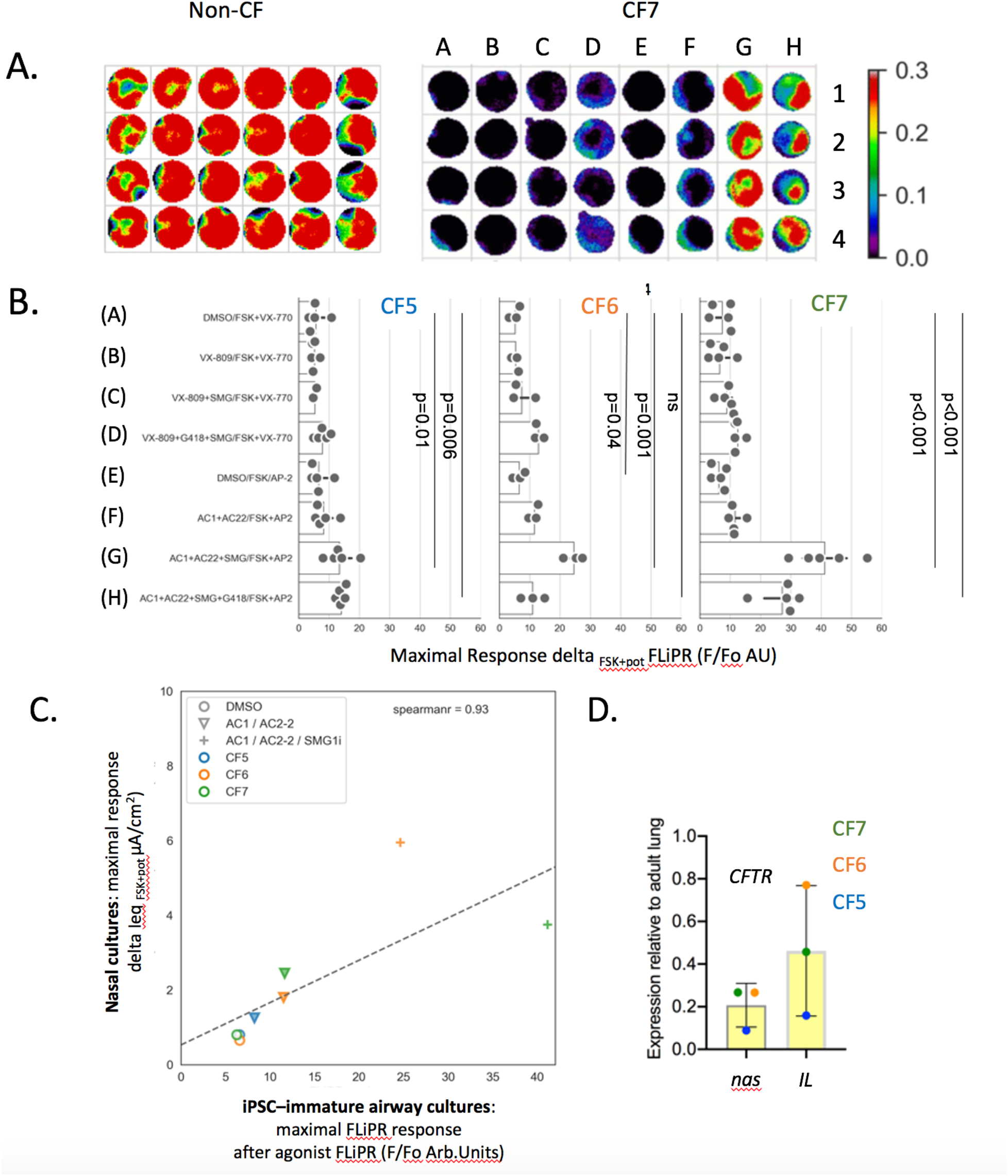
Immature lung cultures generated from iPSCs from three different donors with the nonsense mutation, W1282X, exhibit differential phenotypic responses to modulators. **(A)** Heatmap of peak responses generated from non-CF culture stimulated with 10μM FSK (left) and CF culture (W1282X) treated with combination of small molecules (A-H, concentrations defined in Table 1). **(B)** Bar graph shown the peak response of W1282X-CFTR treated with DMSO (0.1%) or small molecules (defined on y axis with concentrations indicated in **Table 1**). After 5 min baseline, the cells were stimulated with DMSO or FSK +/− 1μM VX-770 (VX) or 1.5μM AP2 (AC). Mean peak responses (of 4 wells in a 96 well plate) after agonist and potentiator shown for each small molecule combination as a solid circle. Three patients homozygous for W1282X were studied (CF5,6 and 7). For CF5, 5 plates generated from 4 differentiations from iPSCs were studied. For CF6, 4 plates from 1 differentiation were studied and for CF7, 5 plates from 2 differentiations were studied. CFTR modulators (VX-809 or AC1+AC2-2), in combination with SMG1i, were effective in increasing the abundance of the truncated W1282X protein in immature lung cultures (**Supp Fig 2A**). **(C)** Correlation plot between FLIPR peak response and Ussing chamber studies (data from (*43*)). **(D)** Basal W1282X-CFTR transcript (left) expression in primary nasal epithelial cultures (nas) and iPSC-derived immature lung cultures (IL) from three donors.

A panel of small molecule interventions was tested for efficacy in rescuing the functional expression of W1282X in three donors, all homozygous for this mutation. These interventions were chosen based on previous experiments in a bronchial cell lines (*8*) and our previous bioelectric studies of primary nasal epithelial cultures (*43*). Based on these previous studies, we expect that only those cultures receiving SMG1i, a small molecule inhibitor of nonsense mediated decay together with protein modulators of the assembly and function (*8*), would enable functional rescue.

As shown in the bar graphs in 4B, the responses observed for each patient-specific culture are reproducible. Interestingly, there were variable responses to the same panel of modulators across the 3 donors. For example, for donor CF5, the magnitude of the response to the modulator combinations is muted relative to the cultures derived from CF6 and CF7. As expected, significant functional rescue was only observed in immature cultures treated with the nonsense mediated decay inhibitor, SMG1i in combination with correctors and potentiator of the truncated W1282-CFTR protein. Interestingly, the premature termination codon (PTC) read-through agent (G418) was not effective augmenting functional rescue of W1282X mediated by SMG1i, as suggested in some but not all studies of primary cultures harboring this nonsense mutation (42). Hence, we show that combinatorial interventions targeting W1282X-CFTR can be tested in patient derived cultures using the FLiPR based high-throughput assay. Further, we show that nonsense mediated decay is limiting modulator activity in these cultures and finally, that different responses are observed for different patient-specific cultures.

As for the studies of F508del-CFTR, we observed a strong correlation (r=0.93) between the patient-specific response to interventions measured by FLiPR in immature lung and responses measured in primary nasal epithelial cultures in the Ussing chamber (data from (*43*) Figure 4C). Unfortunately, due to limitations related to expansion of primary nasal epithelial cultures, the correlation for all conditions, for all three donors, could not be assessed. However, it is clear that the best functional responses in both iPS-derived immature lung and primary nasal epithelial cultures, is achieved after inhibition of nonsense mediated decay using SMG1i.

The mutant CFTR transcript abundance (expressed relative to adult lung, shown in **Fig. 4D**, upper graph), is similar for both the iPSC derived immature lung cultures and the matched nasal cultures.

## DISCUSSION

In these studies, we showed that fluorescence-based channel activity assays of immature lung cultures generated from iPSCs, have the potential to support therapy development for individuals with CF. Such immature lung cultures possess disease-relevant cell types, such as secretory cells, and can be scaled up to enable robust phenotypic screening of therapeutic interventions. As suggested in our proof-of-concept studies, functional responses seen in this novel assay platform recapitulate patient specific responses measured by low-throughput Ussing chamber studies of primary nasal epithelial cultures, thereby supporting the future development of this novel screening platform to aid in precision medicine development for CF individuals currently lacking therapeutic options.

A caveat of our work, relates to the relatively immature airway tissue employed for the phenotypic screen. Although transcriptomic studies show that the submerged, immature lung tissue expresses CFTR and contains CF-relevant cells types including secretory cells such as club and goblet cells (Figure 1), we acknowledge that there will be a deficit of certain other cell types, including ionocytes. Hence, we suggest that the medium to high-throughput iPSC-based screen described here can be used to identify promising compounds that target defects in RNA and protein processing exhibited by CF causing nonsense mutations. Subsequently, compounds identified from these primary screens should be validated in bioelectric assays of fully differentiated primary nasal epithelial tissues as in the current work.

Interestingly, at least for the cultures studied here, there was no difference in the expression of cell type markers between immature airway cultures derived from CF and non-CF iPS cells. These studies support previous publications, suggesting that CFTR mutations do not change cell type distribution (*45*). However, additional transcriptomic studies of lung tissues modeling early lung development for individuals harbouring nonsense mutations, expected to reduce CFTR mRNA by triggering NMD, are required for a more fulsome analysis of the role of CFTR expression on early lung development. Similarly, the impact of CFTR nonsense mutations on proximal airway maturation after transitioning to the air/liquid interface are required in order to test the impact of CFTR expression on airway differentiation.

Although an extremely active area of research, there are no approved treatments targeting the primary defects caused by nonsense mutations in CFTR. Small molecules that facilitate read-through of premature termination codons (PTCs) to enable production of full length CFTR protein are undergoing development. However, only one such compound, called ELOX-2 is currently in clinical trial for subjects carrying the CF-causing nonsense mutation, G542X (*46*), https://www.cff.org/Trials/Pipeline. An alternative approach is to suppress NMD using small molecule inhibitors which has been shown to augment abundance of the truncated W1282X protein in the current work and previous papers (*8, 43, 47*); though such inhibitors may have nonspecific, deleterious effects. Further, such NMD inhibitors will need to be combined with protein modulators in order to correct the primary defects in W1282-CFTR function. Hence, combinatorial treatments aimed at promoting PTC readthrough, inhibition of NMD and modulation of the stability and function of the mutant protein will need to be tested to identify the most effective interventions. We anticipate that the iPS cell phenotypic platform that we describe here will facilitate such urgently needed therapy development by facilitating comparative analysis of multiple therapeutic strategies.

Individuals who are homozygous for nonsense mutations are rare world-wide (188 patients, http://cftr2.org) and as a result, relevant airway tissue models for therapy testing are limited. Immature lung differentiated from iPSC lines are infinitely renewable in principle and we show that after differentiation to an immature tissue, together with the FLiPR assay of CFTR channel activity, they can be employed for testing therapeutic strategies.

Importantly, immature lung cultures from individuals homozygous for F508del exhibited variable phenotypic responses to modulators targeting its mutation class. Likewise, immature lung cultures from individuals homozygous for W1282X also exhibited variable patient-specific responses to small molecules targeting this nonsense mutation. Hence, we provide a resource with which to identify therapeutic targets other than mutant CFTR that may inform companion therapies. Finally, our findings suggest that patient-specific genetic background plays a role in conferring variable drug effect size and highlights the importance of iPS-derived lung cultures for advancing therapy development and precision medicine.

## MATERIALS AND METHODS

### Cell culture: Differentiation to submerged immature lung culture

Human iPS cells were obtained from the Cystic Fibrosis Individualized Therapy (CFIT) program (*48*). The submerged immature lung cultures were generated from iPS cells as previously described (*35*). Human iPS cells were grown on six-well plates (Corning) coated with Matrigel (Corning) and maintained with mTeSR (Stem Cell Technologies). Cultures were expanded weekly with Gentle Cell Dissociation Buffer (GCDR, Stem Cell Technologies) at 70-90% confluency at a 1:10 ratio. For definitive endoderm (DE) induction, single-cell suspensions were generated from five-minute incubation at 37°C followed by scraping and gentle trituration. Cells were plated onto six-well plates in media supplemented with 10 μM Y27632 compound (Stem Cell Technologies) for 24 hours. DE cultures were generated using the StemDiff Definitive Endoderm Kit (Stem Cell Technologies) as per manufacturer’s protocol for 5 days. To differentiate anterior foregut endoderm (AFE) culture, cells were treated with differentiation basal medium (KnockOut DMEM, 10% KnockOut serum replacement, 1% penicillin-streptomycin, 2mM Glutamax, 0.15 mM monothioglycerol, and 1 mM non-essential amino acid) supplemented with FGF2 (500 ng/mL) and SHH (500 ng/mL) for 24 hours. On the second day of AFE differentiation, cells were dissociated into single-cell suspensions and plated onto type IV collagen coated (60 ug/mL, Sigma) 96-well plates at a density of 25,000 cells per well. The media was supplemented with 10 μM Y27632 compound for 24 hours and was changed every 48 hours for an additional three days. For directed differentiation to lung progenitor cells and immature lung cells, cultures were overlayed with differentiation basal medium supplemented with FGF7 (50 ng/mL), FGF10 (50 ng/mL) and BMP4 (5 ng/mL) for 5 days, and then FGF7 (10 ng/mL), FGF10 (10 ng/mL) and FGF18 (10 ng/mL) for 5 days.

### Membrane potential based functional assays

#### Apical Chloride Conductance (ACC) Assay for CFTR function

The ACC assay was used to assess CFTR mediated changed in membrane depolarization using methods as previously described (*40, 49*). In summary, iPSC derived- submerged lung cultures were incubated with zero sodium, chloride and bicarbonate buffer (NMDG 150 mM, Gluconic acid lactone 150 mM, Potassium Gluconate 3 mM, Hepes 10 mM, pH 7.42, 300 mOsm) containing 0.5 mg/ml of FLIPR^®^ dye for 30 mins at 37°C. Wt-CFTR function was measured after acute addition of Fsk (10 μM) or 0.01% DMSO control. Cells were chronically rescued with corrector compounds for 24 hours. Post drug rescue, F508del-CFTR function was measured after acute addition of Fsk (10 μM) and VX-770 (1 μM) or AP-2 (1.5 μM). CFTR functional recordings were measured using the FLIPR^®^ Tetra High-throughput Cellular Screening System (Molecular Devices), which allowed for simultaneous image acquisition of the entire 96 well plate. Images were first collected to establish baseline readings over 5 mins at 30 second intervals. Modulators were then added to stimulate CFTR mediated anion efflux. Post drug addition, CFTR mediated fluorescence changes were monitored and images were collected at 15 second intervals for 70 frames. CFTR channel activity was terminated with addition of Inh172 (10 μM) and fluorescence changes were monitored at 30 second intervals for 25 frames.

### Analysis and heatmap generation

Experiments were exported as multi frame TIFF images of which every frame recorded the entire plate. Pixels outside of well areas were filtered out using the initial signal intensities and wells containing opened organoids were separated. All traces were normalized to the last point of the baseline intensity. Peak response for each pixel was calculated as the maximum deviation from baseline. During the stimulation segment, fluorescence intensity increased for CFTR function. Heatmap representation was generated from the peak response of each pixel and the mean response trace of wells was generated by averaging the corresponding pixel traces.

### Real-time Quantitative PCR

As previously described (*21*), total mRNA was extracted using RNeasy^®^ Plus Micro Kit, following enclosed instructions. After measuring the spectrophotometric quality of extracted RNA through 260/280 ratios of 2.0 and 260/230 ratios of 1.8-2.2, mRNA samples used to reverse transcribe 1 μg of cDNA using iScript™ cDNA Synthesis Kit. Quantitative real-time PCR was performed with PowerUP SYBR Green Mastermix Master Mix on ViiA7 (Applied Biosystems). Gene expression is normalized to house-keeping gene GAPDH and expressed relative to control human tissue RNA extracts (2^ΔΔCT). A total run of 40 cycles. Cycle threshold (CT) values above 38 were considered “not expressed”. The primers used for amplification are described in the following table.

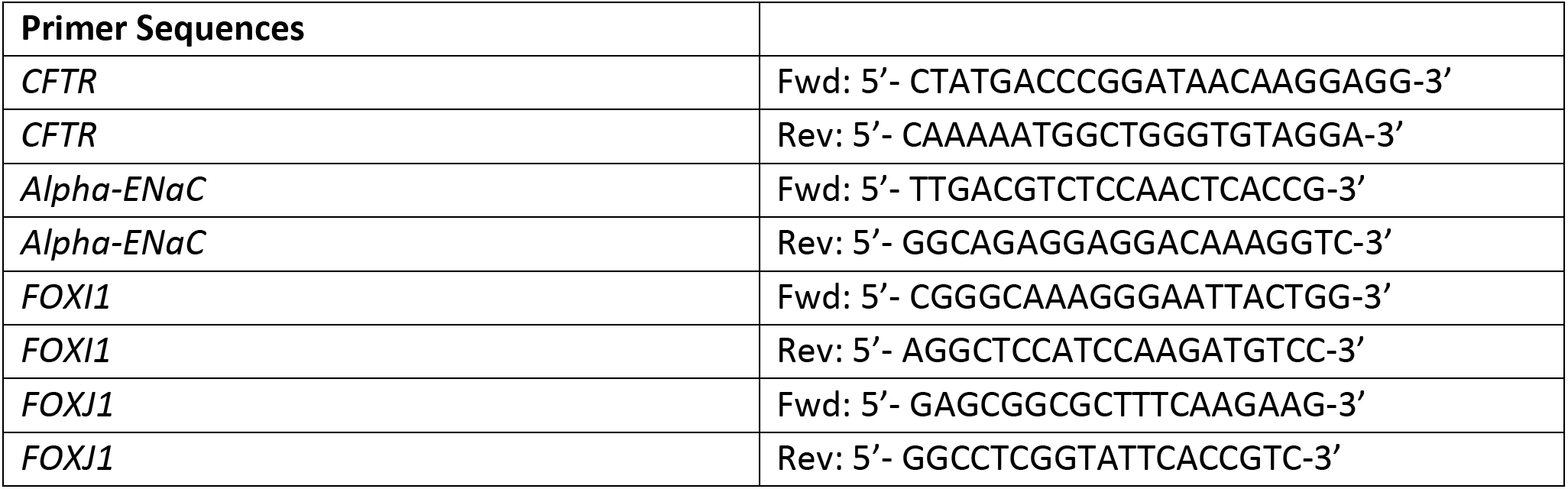
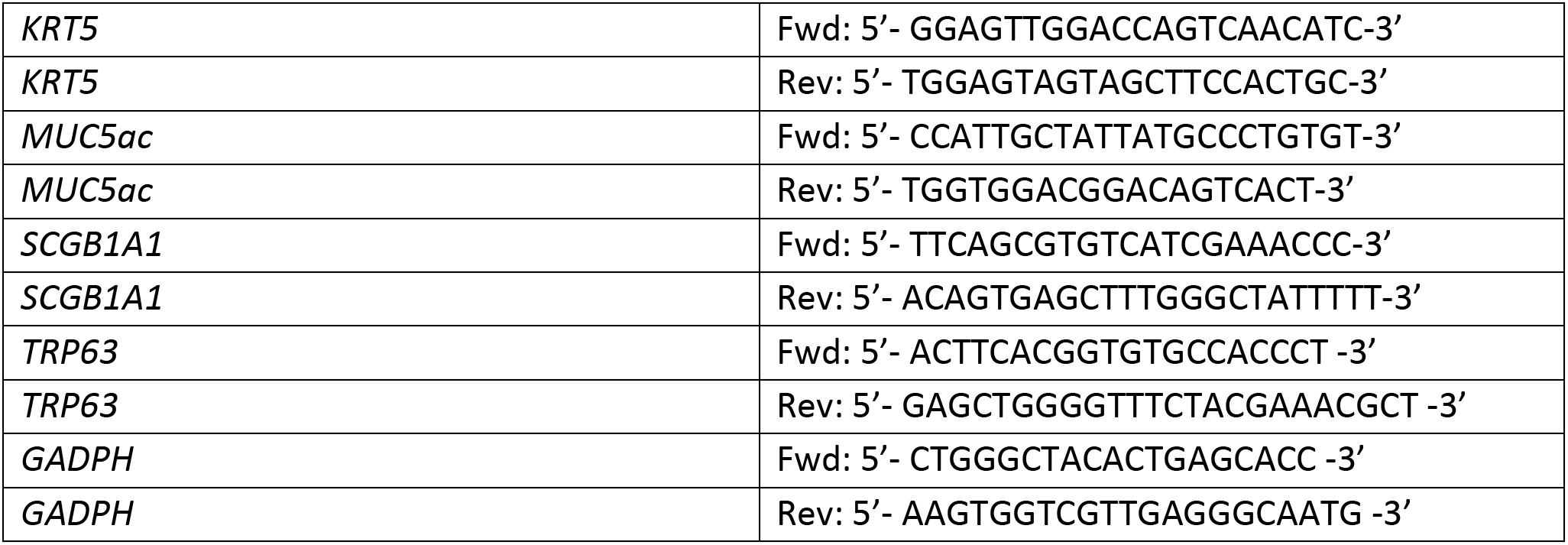

### RNA Sequencing and Analysis

RNA samples were extracted using methods described previously (*50, 51*). RNA samples with an RNA integrity number (RIN) greater than 8.5 was submitted to The Centre of Applied Genomics (TCAG) at SickKids for bulk RNA sequencing. In brief, RNA libraries were generated using NEB Ultra II Directional mRNA with an average of 69,954,038 reads from each library on the Illumina HiSeq 2500 platform using high-throughput V4 flowcells.

Post sequencing quality control was performed using open-source software FastQC (https://www.bioinformatics.babraham.ac.uk/projects/fastqc/). Trim galore (*52*) was used to remove low quality sequences and trim adapters. Paired-end reads were aligned to the human reference genome (hg38) using STAR (version 2.7.1a) (*53*). The resulting bam files containing aligned sequences were subsequently processed using SAMtools (*54*), and raw counts generated with featureCounts (*55*) were used for downstream analysis (transcripts per gene). R package DESeq2 (v.1.24.0) (*56*) was used to calculate size factors for each sample and perform regularized-logarithm rlog transformation of read counts. The 100 genes with the highest variance in expression across all samples were subjected to principal component analysis (PCA).

### Immunofluorescence

Samples were fixed in 4% paraformaldehyde and then washed three times with PBS, 5 mins per wash at room temperature. Cell permeabilization was performed using 0.05% TritonX-100 followed by three PBS washes. Samples were blocked using 5% BSA for 1 hour and incubated with primary antibody against TTF1 (NKX2-1) or EPCAM overnight. After removal of primary antibody, samples were washed 3 times with PBS, 5 mins per wash and incubated with secondary antibodies and nuclear marker DAPI for 1 hour. Samples were then washed 3 times with PBS, 5 mins per wash at room temperature. Images were acquired on the SP8/STED confocal microscope (Leica).

### Western blotting

Samples were collected in ice cold PBS and pelleted through centrifugation at 4°C (500*g* for 7 mins). Post centrifugation, the cell pellet was re-suspended in 200μL of modified radioimmunoprecipitation assay butter (50 mM Tris-HCl, 150 mM NaCl, 1 mM EDTA, pH 7.4, 0.2% (v/v) SDS and 0.1% (v/v) Triton X-100) containing a protease inhibitor cocktail for 10 min. After centrifugation at 13,000 rpm for 5 min, the soluble fractions were analyzed by SDS-PAGE on 6% Tris-Glycine gel. After electrophoresis, proteins were transferred to nitrocellulose membranes and incubated in 5% milk and CFTR bands were detected using the mAb 596. Calnexin (CNX) was used as a loading control and detected using a Calnexin-specific rAb (1:5000). The blots were developed with using the Li-Cor Odyssey Fc (LI-COR Biosciences, Lincoln, NE, USA) in a linear rage of exposure (1-20 min). Relative levels of CFTR protein were quantitated by densitometry of immunoblots using ImageStudioLite (LI-COR Biosciences, Lincoln, NE, USA) (*57, 58*).

## Supplementary Materials

**Supplementary Figure 1:** Representative F508del-CFTR protein expression in lung submerge after 48h pre-treatment with DMSO (0.1%), 3μM VX-809, 0.5μM AC1 + 3μM AC2-1 or 0.5μM AC1 + 3μM AC2-2.

**Supplementary Figure 2:** Correction of W1282X mutation by CRISPR-Cas9 editing, confers CFTR channel activity in submerged, immature lung cultures from donor #4. CFTR channel activity was measured using the FLiPR assay and the bars represent mean +/− SD in 4 technical replicates.

**Table 1: Concentrations employed for small molecules**

## ACKNOWLEDGMENTS

iPS cells and primary nasal cell cultures were obtained through the CF Canada-SickKids Program for Individualized CF Therapy (CFIT). The SMG1i was obtained through Cystic Fibrosis Foundation Therapeutics and the small molecules AC1, AC2-1, AC2-2 and AP2 were provided by AbbVie. We thank Ashvani Singh for reviewing this manuscript.

## Author contributions

Jia Xin Jiang: designed and performed experiments, revised the manuscript

Leigh Wellhauser: performed experiments

Onofrio Laselva: performed experiments and revised the manuscript

Irina Utkina: Data analysis

Zoltan Bozoky: Data analysis

Tarini Gunawardena: Performed experiments

Zoe Ngan: Performed experiments

Sunny Xia: Data collection

Paul D.W. Eckford: Documentation oversight, reviewed and edited the manuscript

Felix Ratjen: Provided scientific insight, reviewed and edited the manuscript

Theo Moraes: Provided scientific insight, reviewed and edited the manuscript

John Parkinson: Data analysis and interpretation, reviewed the manuscript

Amy P. Wong: Data analysis and interpretation, reviewed the manuscript

Christine E. Bear: Conceived and designed the work, drafted and revised the manuscript

## Data and materials availability

Raw sequence data is available through the NCBI sequence read archive (https://www.ncbi.nlm.nih.gov/sra) with the BioProject ID PRJNA721455.

