## supplemental file for "A new platform for high-throughput therapy testing on iPSC-derived, immature airway from Cystic Fibrosis Patients"

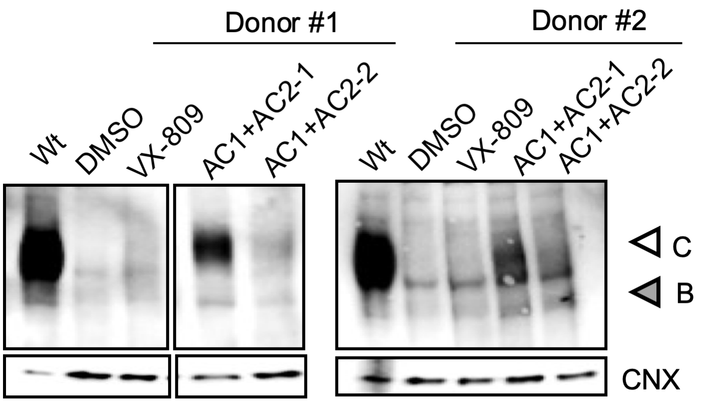


**Fig. S1:** Representative F508del-CFTR protein expression in immature airway epithelium after 48h pre-treatment with DMSO (0.1%), 3µM VX-809, 0.5µM AC1 + 3µM AC2-1 or 0.5µM AC1 + 3µM AC2-2. C: mature, complex-glycosylated CFTR; B: immature, core-glycosylated CFTR; CNX, Calnexin as loading control.

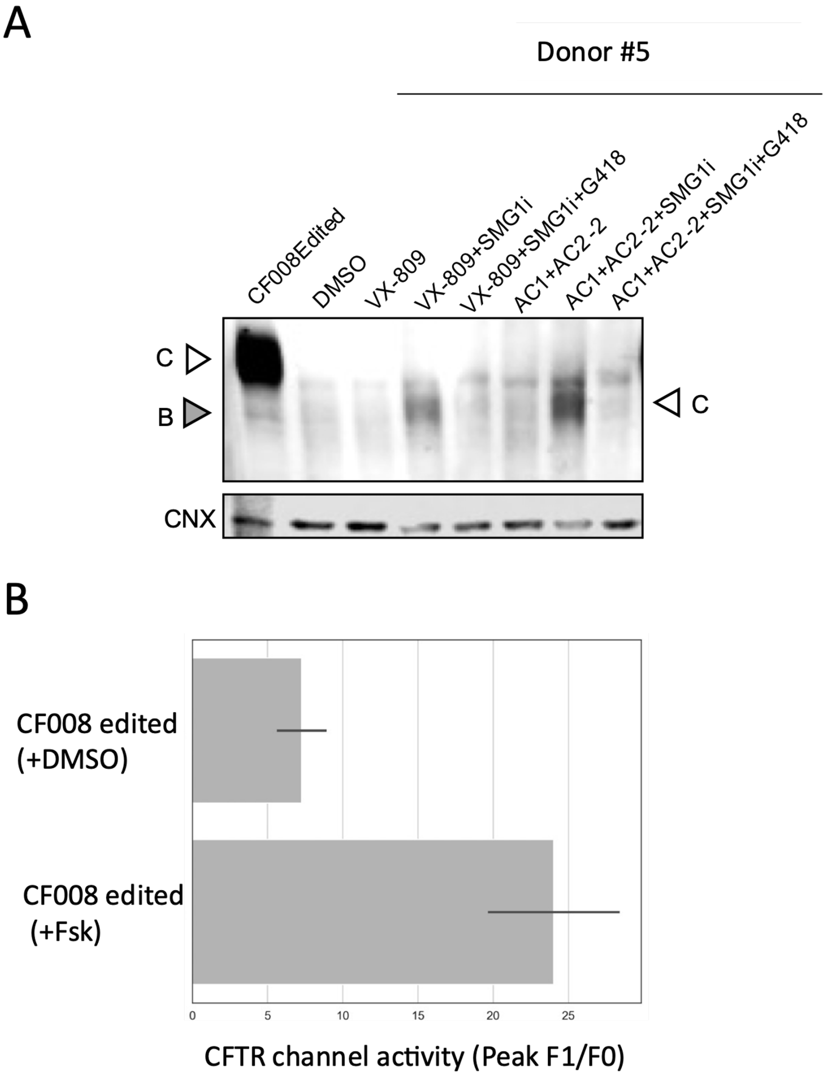


**Fig. S2: Correction of W1282X mutation confers CFTR channel activity in submerged, immature lung cultures from donor=CF008**. The iPSCs line from a donor, homozygous for W1282X, was corrected on one allele. (**A**) Representative W1282X-CFTR protein expression in lung submerge after 48h pre-treatment with DMSO (0.1%), 3µM VX-809, 3µM VX-809+ 0.5µM SMG1i, 3µM VX-809+ 0.5µM SMG1i+ 200µg/mL G418, 0.5µM AC1 + 3µM AC2-2, 0.5µM AC1 + 3µM AC2-2+ 0.5µM SMG1i or 0.5µM AC1 + 3µM AC2-2+ 0.5µM SMG1i+200µg/mL G418. (**B**) 10µM Forskolin activated CFTR channel function was conferred with correction in the differentiated to immature lung cultures. CFTR channel activity was measured using the FLiPR assay and the bars represent mean-/+ SD in 4 technical replicates.

**Table 1**

| Small molecules | Concentration |
|---|---|
| VX-809 | 3 µM |
| AC-1 | 0.5 µM |
| AC2-2 | 3 µM |
| SMG1i | 0.5 µM |
| G418 | 200 µg/mL |
